# Acute depletion of METTL3 identifies a role for *N*^*6*^-methyladenosine in alternative intron/exon inclusion in the nascent transcriptome

**DOI:** 10.1101/2020.09.10.291179

**Authors:** Guifeng Wei, Mafalda Almeida, Greta Pintacuda, Heather Coker, Joseph S Bowness, Jernej Ule, Neil Brockdorff

## Abstract

RNA *N*^*6*^-methyladenosine (m6A) modification plays important roles in multiple aspects of RNA regulation. m6A is installed co-transcriptionally by the METTL3/14 complex, but its direct roles in RNA processing remain unclear. Here we investigate the presence of m6A in nascent RNA of mouse embryonic stem cells (mESCs). We find that around 10% m6A peaks are in introns, often close to 5’-splice sites. RNA m6A peaks significantly overlap with RBM15 RNA binding sites and the histone modification H3K36me3. Interestingly, acute dTAG depletion of METTL3 reveals that inclusion of m6A-bearing alternative introns/exons in the nascent transcriptome is disrupted. For terminal or variable-length exons, m6A peaks are generally located upstream of a repressed 5’-splice site, and downstream of an enhanced 5’-splice site. Intriguingly, genes with the most immediate effects on splicing include several components of the m6A pathway, suggesting an autoregulatory function. Our findings demonstrate a direct crosstalk between m6A machinery and the regulation of RNA processing.

## Introduction

RNA is subject to diverse post-transcriptional modifications that have emerged as new layers of gene regulation (Fu et al., 2014; Roundtree et al., 2017a; Yue et al., 2015). Among these, *N*^*6*^-methyladenosine (m6A) is the most prevalent and abundant internal RNA modification on mRNA. m6A was initially identified in the 1970s (Desrosiers et al., 1974; Perry et al., 1975), and the enzyme that catalyzes this modification was described in the mid-1990s (Bokar et al., 1994; Bokar et al., 1997). Accumulating evidence suggests that RNA m6A modifications are in most part installed by the METTL3/14 core heterodimer (Liu et al., 2014), which together with accessory proteins WTAP (Ping et al., 2014), VIRMA (Schwartz et al., 2014), RBM15/15B (Patil et al., 2016), CBLL1 (Ruzicka et al., 2017), and ZC3H13 (Knuckles et al., 2018; Wen et al., 2018) forms the m6A writer complex. Structural studies have revealed that METTL3 is the only catalytic subunit, while METTL14 has a degenerate active site and maintains the complex’ integrity and substrate RNA recognition (Sledz and Jinek, 2016; Wang et al., 2016a; Wang et al., 2016b). Similar to DNA and histone modifications pathways, the m6A pathway has specific eraser (FTO and ALKBH5) and reader proteins (YTH-domain-containing proteins, YTHDC1/2 and YTHDF1/2/3) (Zaccara et al., 2019).

Global m6A patterns have been profiled using m6A specific antibodies coupled to high-throughput sequencing (Dominissini et al., 2013; Ke et al., 2015; Linder et al., 2015; Meyer et al., 2012). Antibody-free m6A profiling methods, MAZTER-seq (Garcia-Campos et al., 2019) and m6A-REF-seq (Zhang et al., 2019) have been developed since, but are limited to a subset of the m6A (m6ACA) sites. Extensive m6A profiling in a variety of RNA populations from diverse species and tissues have revealed that the majority of mRNAs are m6A modified with preferred sites occurring in clusters, most commonly in the 3’ UTR and around the stop codon (Dominissini et al., 2012; Meyer et al., 2012). Individual m6A sites have the consensus sequence DRACH (Dominissini et al., 2012; Linder et al., 2015; Meyer et al., 2012). The installation of m6A by the writer complex occurs co-transcriptionally and sites are found both in exons (the majority) and introns (Ke et al., 2017; Louloupi et al., 2018). An important factor for targeting m6A to defined sites is the RNA binding protein RBM15/15B, a subunit of the m6A writer complex (Coker et al., 2020; Patil et al., 2016). Additionally, the METTL14 subunit recognizes the histone modification H3K36me3, which is enriched within gene bodies of active genes (Huang et al., 2019). Finally some transcription factors (TF) have been proposed to facilitate m6A targeting, for example SMAD2/3 (Bertero et al., 2018) and CEBPZ (Barbieri et al., 2017), although only for a small number of transcripts in certain conditions and/or cell types.

The m6A modification has important functions in mRNA metabolism, notably in the regulation of RNA processing (Alarcon et al., 2015b), nuclear export (Roundtree et al., 2017b), turnover (Ke et al., 2017; Liu et al., 2020; Wang et al., 2014), and translation (Barbieri et al., 2017; Wang et al., 2015). There are however contradictory findings, for example in relation to alternative splicing (Alarcon et al., 2015a; Ke et al., 2017; Xiao et al., 2016), translation (Wang et al., 2015; Zaccara and Jaffrey, 2020), and X chromosome inactivation (Coker et al., 2020; Nesterova et al., 2019; Patil et al., 2016). Confounding factors include the difficulty in discriminating primary and secondary effects following chronic long-term knockout/knockdown of m6A writers/readers, and cell lethality effects linked to the important role of m6A in essential cell functions (Barbieri et al., 2017; Xiao et al., 2018).

In this study we analyse m6A in the nascent transcriptome of mouse embryonic stem cells (mESCs) with a view to uncover important roles of METTL3 complex in alternative splicing. We make use of the dTAG protein degron system (Nabet et al., 2018) to rapidly and acutely deplete the m6A methyltransferase METTL3, thereby discriminating primary from secondary effects. We characterize intronic m6A sites and show that they correlate with both RBM15 binding and the histone H3K36me3 modification. Functional analysis following acute depletion of METTL3 uncovered numerous changes to alternative splicing occurring at or near to m6A methylation sites. Intriguingly, m6A-dependent alternative splicing affects several genes encoding factors in the m6A pathway, suggestive of autoregulatory functions.

## Results

### Mapping m6A in the nascent mESC transcriptome

Chromatin associated RNA (ChrRNA) is substantially enriched for nascent transcripts (Nesterova et al., 2019). Thus, to investigate the roles of m6A in nascent RNA processing in mouse embryonic stem cells (mESCs), we performed MeRIP-seq from ChrRNA, referred to henceforth as ChrMeRIP (Figure 1A and S1A). Sequencing of input showed that 70∼80% of reads are intronic (Figure S1B). To minimize specific antibody bias, we used two commercially available m6A antibodies (SySy and Abcam) to identify high confidence m6A-modified RNA regions. Using maximum ORF and longest ncRNA isoforms as representative transcripts (see Methods), refined peak calling analysis (see Methods) classified 5277, 5472, and 6319 m6A peaks into Confidence group 1 (high), Confidence group2 (medium), and Confidence group3 (low), respectively (Figure 1B-D and S1C,D). The trend of m6A peak intensity in the different groups accords with their confidence classifications (Figure 1B and S1E). Notably, the overlap between peaks from SySy and Abcam antibodies is approximately half, which is similar to the differences in peak detection surveyed between studies (Figure 1C) (McIntyre et al., 2020). Despite the large fraction of intronic reads in the input, only 6.2% of confidence group1 (Cfg1) and 10.3% of confidence group2 (Cfg2) peaks are from intronic regions (Figure 1D and S2A), as defined by the position of the single-nucleotide peak summit (see Methods). This is slightly higher than previously reported for MeRIP-seq or m6A-CLIP studies using only messenger RNA from mESCs (Figure S2B) (Batista et al., 2014; Geula et al., 2015; Ke et al., 2017), and is in line with ChrMeRIP-seq from HeLa cells (Ke et al., 2017). The majority of intronic m6A modification occurs in protein-coding genes rather than noncoding RNAs (Figure 1D and S2A,B).

**Figure 1.**
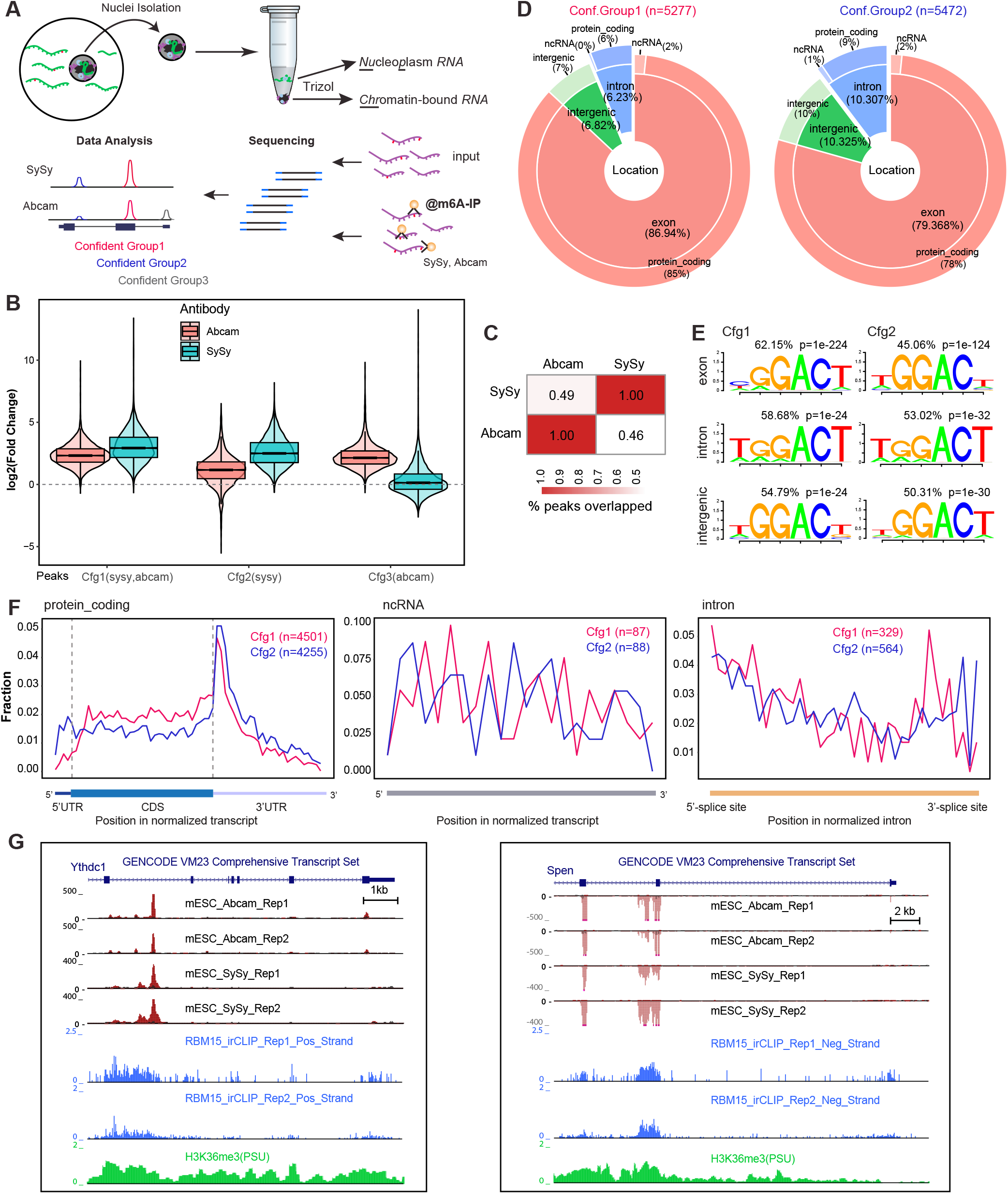
ChrMeRIP-seq reveals that 6-10% of m6A peaks are located in introns. (A) Schematic illustrating the experimental and computational workflow for ChrMeRIP-seq. Chromatin-associated RNAs (ChrRNA) enriched for introns was used for MeRIP with two commercially available m6A antibodies (SySy and Abcam). Three confidence groups of m6A sites were identified. (B) Boxplots showing the m6A intensity distributions for Confidence group 1 (Cfg1), Confidence group 2 (Cfg2), and Confidence group 3 (Cfg3) groups (see Methods for details). Pink and cyan represent m6A intensity from Abcam and SySy antibodies respectively. (C) Heatmap showing the peak overlap between two antibodies. (D) Pie chart output from RNAmpp analysis showing the distribution of m6A peaks for Cfg1 (left) and Cfg2 (right) group. Peak numbers are indicated above. The MaxORF and longestNcRNA isoform was chosen for each gene. (E) Most representative motifs called for each subgroup (exonic, intronic, and intergenic) in Cfg1 and Cfg2 group. (F) RNA Metaprofiles (RNAmpp) of m6A peaks in transcriptome for Cfg1 (red) and Cfg2 (blue) groups. Left plot is for aggregated protein-coding gene, middle for noncoding RNA, and right for normalized intron. (G) UCSC genome browser screenshots showing example genes (left, Ythdc1 intron11; right Spen intron2) harboring intronic m6A methylation. From top to bottom, tracks denote gene annotation, MeRIP-seq (Abcam 2 replicates, SySy 2 replicates), RBM15 irCLIP-seq (2 replicates), and H3K36me3 ChIP-seq.

We developed RNA metaprofile plot (RNAmpp) to describe the distribution of m6A in the nascent transcriptome (see Methods). m6A peaks from both confidence groups 1 and 2 are enriched around stop-codon regions or at the beginning of the 3’-UTR in mRNAs, and at all genomic locations the canonical DRACH m6A motif (GGACU) is the most highly represented motif (Figure 1E,F and S2C). The few m6A peaks which map to lncRNAs are not close to the 3’-end regions, but are rather distributed randomly across the transcripts, as exemplified by *Xist, Norad*, and *Malat1* lncRNA genes (Figure S2D) (Coker et al., 2019; Patil et al., 2016). We found several clear examples of intronic m6A peaks located close to 5’-splice sites, such as the two intronic m6A peaks from *Ythdc1* intron 11 and *Spen* intron 2 (Figure 1G).

Given that the pattern of exonic m6A has been extensively characterized (Dominissini et al., 2012; Meyer et al., 2012), we sought to specifically investigate the deposition pattern and characteristics of intronic m6A modification. To reduce bias, we compared the GC content, conservation level, and relative position of intronic m6A peaks to their size-matched control regions derived from random regions within the same intron (see Methods). This analysis shows that intronic m6A-methylations are more common in regions which have high GC content, are evolutionarily conserved, and are in proximity to 5’-splice sites (Figure 2A-D and S3A-C). When compared to randomly-chosen introns from the same genes as control, longer introns are preferentially methylated (Figure S3D).

**Figure 2.**
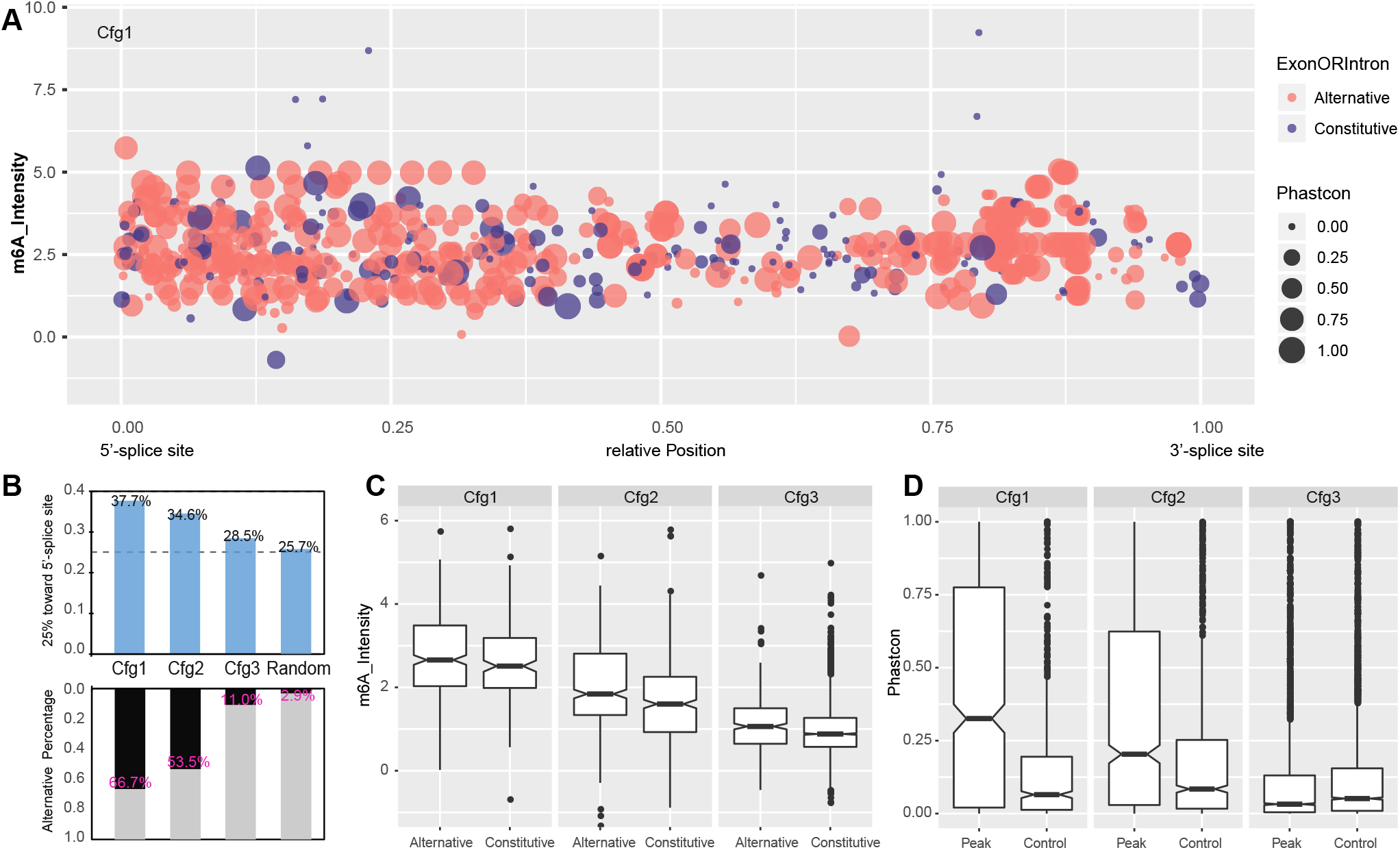
Pattern of intronic m6A modification. (A) Dot plot show the relative position, m6A intensity, conservation, and location in alternative or constitute introns for all intronic m6A methylation in Cfg1. Here 0 and 1 in the X-axis represent the 5’- and 3’-splice sites respectively. The Y-axis denotes m6A intensity calculated as an average of all replicates. Dot area indicates the PhastCon conversation score from UCSC browser. Red and blue dot color denotes location in alternative and constitutive introns respectively. (B) Barplots (top) showing the fraction of m6A peak located in first quarter (close to 5’-splice site) of introns for Cfg1, Cfg2, and Cfg3 classes, as well as random simulated peak summits. Barplots (bottom) showing the fraction of m6A peaks located in alternative exon/introns for all the groups. The percentages for each bar were labelled. (C) Boxplots showing intensity of intronic m6A peaks located in alternative and constitute intron for all classes. (D) Boxplot of PhastCon scores for all classes of m6A peaks located in intron regions annotated from the MaxORF_LongestNcRNA isoforms, compared with controls matched for size and intron-of-origin.

Most exonic methylation is deposited around stop-codon regions, as previously noted (Dominissini et al., 2012; Meyer et al., 2012), but intronic methylation sites are fairly evenly distributed relative to host transcripts (Figure S3E). Therefore, we queried in particular whether intronic methylated regions are located close to alternative exons or introns using the comprehensive GENCODE annotation set (vM24). Indeed, we found that higher confidence groups of m6A-methylation were more likely to reside in alternative intron/exon regions (see Methods) than low confidence groups or random genomic regions (Figure 2A,B). This indicated a potential role of METTL3 complex and its methylation sites in the regulation of alternative splicing.

### Intronic m6A modifications correlate with RBM15 binding and H3K36me3

The RNA-binding protein RBM15 plays an important role in targeting m6A to defined sites in mRNA (Patil et al., 2016). We went on to examine if this pathway is linked to the intronic m6A sites that we observe in nascent RNA. To map binding sites for RBM15 in mESCs we performed infrared crosslinking immunoprecipitation followed by sequencing (irCLIP-seq) (Zarnegar et al., 2016), making use of an mESC line in which both emGFP-Rbm15 and Xist RNA are induced by treatment with doxycycline (Coker et al., 2020) (Figure S4A). RBM15 interacts with the Xist A-repeat region and contributes to the deposition of m6A methylation at sites immediately downstream (Coker et al., 2020; Patil et al., 2016) (Figure S4B), and thus provides a useful positive control.

Cross-linking induced truncation sites (CITS or RT stops) are the main signature occurring in irCLIP-seq datasets. The RNA meta-profile plot shows where these CITSs reside across normalized transcripts, with two main peaks at the start of the transcript and near the stop-codon region (Figure 3A,B), in agreement with the RBM15/15B binding profile in human cells (Patil et al., 2016). Examination of Rbm15 binding across introns shows a preference for the 5’-splice site, consistent with the profile of intronic m6A (Figure 3B and S4C). Motif analysis of CITSs revealed an RBM15 binding consensus comprising 3 or 4 consecutive U bases, both for exonic and intronic sites (Figure 3C and S4D). This is also the case for crosslinking-induced mutations (CIMS) (Figure 3C). Importantly, we found that RBM15 binding is centered at m6A peak summits for all exonic, intronic, and intergenic regions and in general correlates with peak confidence (Figure 3D and S5A,B). We further split each m6A confidence group by strong or weak RBM15 binding, based on whether strong RBM15 biding sites (CITS>=3 in 2 replicates) intersected the m6A peak (Figure 3E-J and S5C-E). Over half of sites in all sub-groupings of Cfg1 and Cfg2 were assigned as strong RBM15-binding, with the only exceptions being Cfg3 intronic and intergenic groups (Figure 3E,H, and S5C). RBM15 binding was centered on m6A peaks across all the confidence groups (Figure 3F,I and S5D).

**Figure 3.**
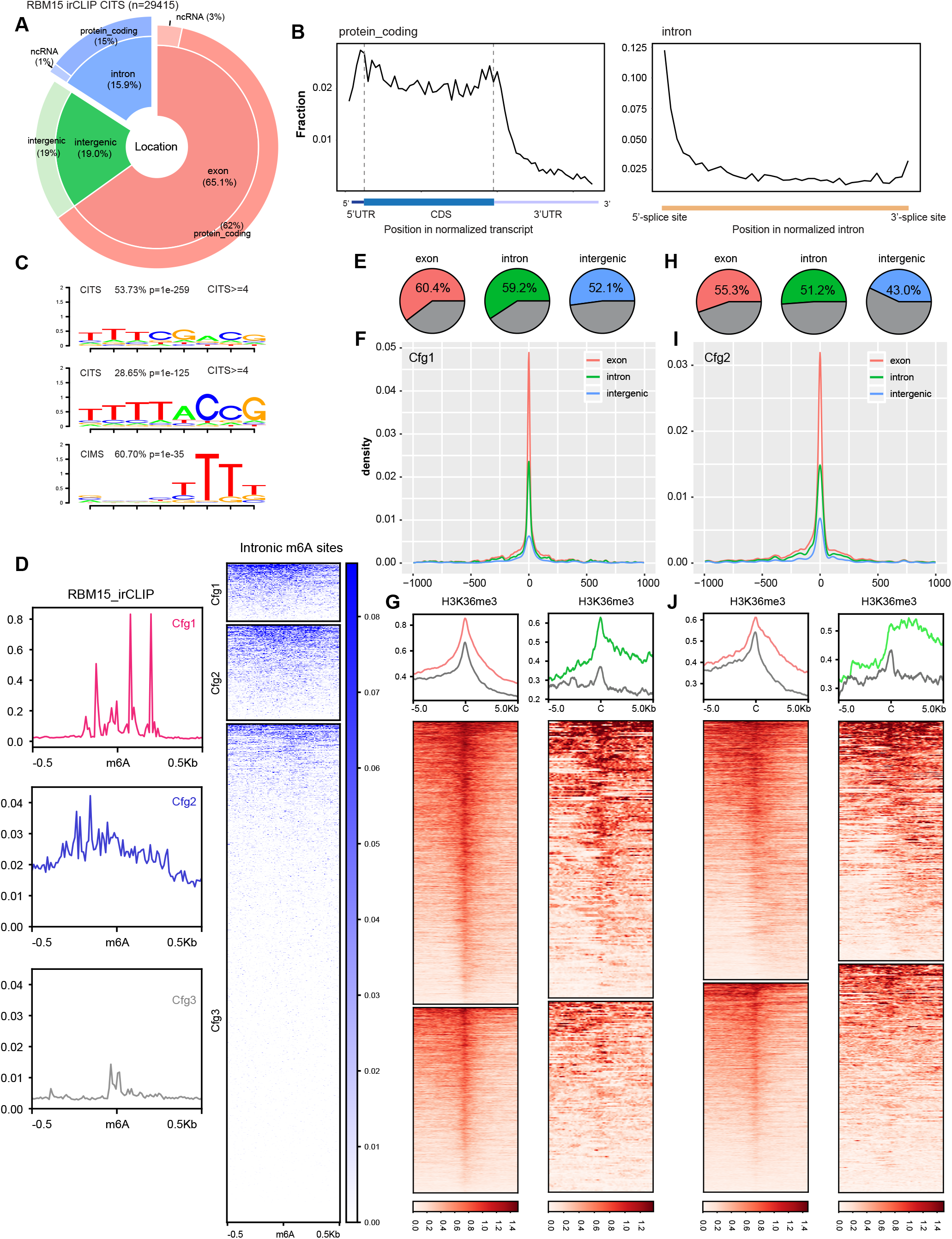
Intronic m6A methylation correlates with RBM15 binding and H3K36me3. (A) Pie chart shows the distribution of RBM15 binding sites, calculated from irCLIP cross-linking induced truncation sites (CITS >=3). (B) The RNA binding profiles of RBM15 binding in the transcriptome, calculated for aggregated gene models for protein-coding genes (left) and introns (right) shown below. (C) RNA-binding motifs occurring at RBM15 CITS and CIMS. (D) The RBM15 binding (CITS) metaprofile and heatmap for intronic m6A peaks of 3 different confidence groups (red, blue, and grey for Cfg1, Cfg2, and Cfg3 respectively). The color key is shown on the right. 0.5kb strand-specific flanking regions on each side of m6A peak summit are included for plot. (E) Pie charts illustrating the fraction of Cfg1 m6A peaks with strong RBM15 CITS (>=3) sites within 1kb flanking regions. Exonic, intronic, and intergenic m6A peaks are shown from left to right. (F) The RBM15 binding site distribution (CITS>=3) centered on m6A peaks. Red, green, and blue lines represent exonic, intronic, and intergenic m6A peaks from Cfg1. (G) Metaprofiles and heatmaps of H3K36me3 for Cfg1 m6A peaks with strong (red and green) and weak (grey) RBM15 binding. Red and green curves denote exonic and intronic m6A peaks respectively. 5kb regions flanking each m6A peak summit are included for the heatmaps, with color keys of H3K36me3 shown below. (H-J) Same as (E-G) but for Cfg2 m6A peaks.

In addition to RBM15, the histone modification H3K36me3 has been proposed to play a role in directing m6A to defined sites in mRNA, mediating interaction with the METTL14 subunit of the core m6A complex (Huang et al., 2019). Consistent with this model we observe that high confidence exonic and intronic m6A peaks correlate with maximal H3K36me3 density (Figure S5F). Furthermore, m6A peaks with strong RBM15 binding also reside within higher H3K36me3 density for both exonic and intronic sites (Figure 3G,J and S5E). Taken together, our observations indicate that intronic m6A sites show equivalent correlations with both RBM15 binding and H3K36me3 density to those seen for exonic sites, suggesting that similar targeting mechanisms function in both contexts.

### Rapid depletion of METTL3 using the dTAG system

Functional analysis of the METTL3/14 m6A writer complex using gene knockout/knockdown has provided conflicting results in terms of the importance of m6A in regulating splicing. A confounding factor is that m6A has roles in mRNA stability and translation (Fu et al., 2014), and it is challenging to discriminate primary and secondary effects resulting from chronic or incomplete loss of function. To address this, we developed an acute Mettl3 knockout model using the dTAG degron system (Nabet et al., 2018). FKBP12^F36V^ was fused in-frame into the C-terminus of Mettl3 in XX female mESCs with doxycycline-inducible Xist using CRISPR-Cas9-mediated knock-in (Figure 4A). In two independent clones (C3 and H5), the level of METTL3_FKBP12^F36V^ expression was very similar to that of endogenous METTL3 and the dTAG-13 treatment did not affect subcellular localization. We also confirmed that the FKBP12^F36V^ insertion does not interrupt the level of the neighboring TOX4 protein, whose 3’-UTR overlaps with last two coding exons of *Mettl3* in an antisense manner (Figure 4A,C). Following treatment with dTAG-13, METTL3_FKBP12^F36V^ protein levels were rapidly depleted, within 30min (Figure 4B). We also observed a strong reduction in levels of METTL14, which forms a stable heterodimer with METTL3, suggesting heterodimer formation is important for METTL14 protein stability (Figure 4B,C and S6A).

**Figure 4.**
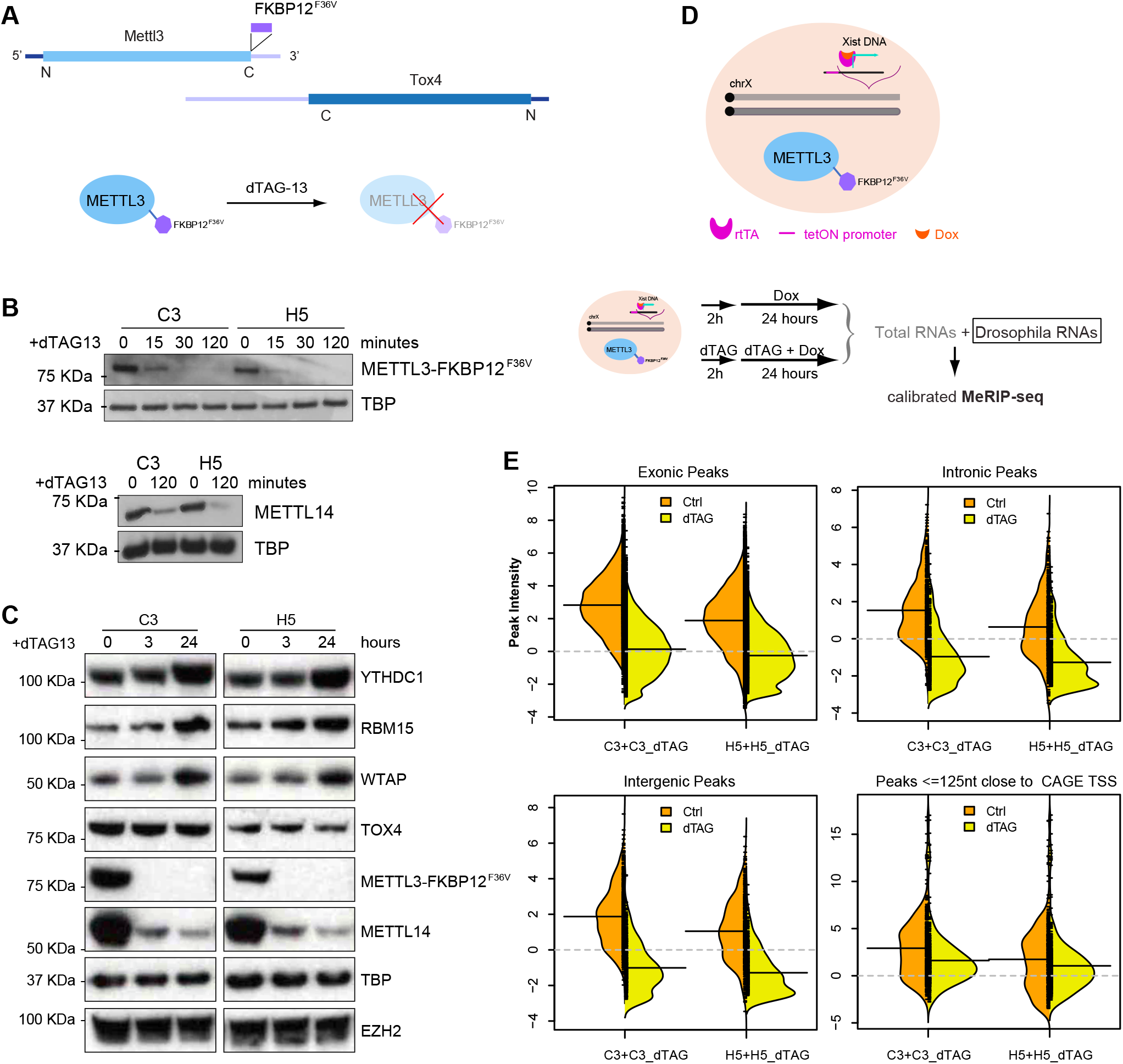
Acute depletion of METTL3 by dTAG system. (A) Schematic illustrates the FKBP12^F36V^ insertion into the stop codon of *Mettl3* gene, which overlaps with Tox4 in an antisense manner. dTAG-13 molecules engage FKBP12^F36V^ to trigger degradation of the fusion protein. (B) Western blots show degradation of METTL3_FKBP12^F36V^ in a dTAG time-course treatment experiment (15min, 30min, 120min) in two independent clones C3 and H5 (top). Lower panel shows METTL14 protein levels in C3 and H5 clones upon 120min dTAG-13 treatment. TBP acts as the loading control. (C) Western blots show the protein levels for YTHDC1, RBM15, WTAP, TOX4, together with METTL3_FKBP12^F36V^ and METTL14 in C3 and H5 clones upon 3 hours and 24 hours dTAG-13 treatment. TBP and non-m6A-modified gene EZH2 as loading control. (D) Schematic showing FKBP12^F36V^ inserted into the Mettl3 locus of hybrid XX mESCs expressing doxycycline-inducible Xist, and the calibrated MeRIP-seq workflow with Drosophila RNAs as a spike in (Bottom). Xist was induced after 2 hours dTAG-13 treatment. (E) Beanplot of the calibrated m6A intensity distributions for peaks classified as exonic, intronic, intergenic from SySy antibody, as well as peaks within 125nt of CAGE TSS. Left and right beanplots show clones C3 and H5 respectively. Orange and yellow back-to-back plots represent Ctrl and dTAG respectively. The black solid line denotes the mean of each distribution and grey dashed line represents the threshold of non-enrichment.

We next sought to examine global dependence of m6A on the METTL3 complex by performing calibrated MeRIP-seq (see Methods) after 26 hrs dTAG-13 treatment. Xist RNA, a useful indicator for m6A deposition (Coker et al., 2020; Ke et al., 2015; Linder et al., 2015; Nesterova et al., 2019; Patil et al., 2016), was induced after 2 hrs dTAG-13 treatment (Figure 4D). For untreated cells, m6A peaks were found at previously annotated sites including Xist RNA (Nesterova et al., 2019), indicating that METTL3_FKBP12^F36V^ retains functionality for m6A modification deposition (Figure 4E and S6B,C). Following dTAG-13 treatment, most Xist m6A peaks were undetectable, including the characteristic sites downstream of the Xist E-repeat (Figure S6B). Moreover, the majority of exonic, intronic, and intergenic m6A peaks became indistinguishable in intensity from input (Figure 4E and S6C). The only exception were peaks that are close to transcript start sites (TSSs), including in Xist RNA, which likely represent m6Am modification (Akichika et al., 2019) that is installed by PCIF1 rather than METTL3/14 (Figure 4E and S6B,C). Collectively, these results demonstrate that the dTAG system enables rapid acute depletion of METTL3 and m6A in mRNA.

### Acute depletion of METTL3 reveals a role for m6A in alternative splicing

We went on to examine the effect of METTL3 depletion on splicing by analysing the newly synthesized transcriptome using 4sU-seq after dTAG-13 treatment for 3 hrs, followed by a short 4sU incorporation (30 min) (Figure 5A). This enabled us to explore transcriptional or co-transcriptional changes upon depletion of METTL3 whilst limiting indirect effects from m6A-mediated RNA destabilization (Liu et al., 2020; Wang et al., 2014). We validated that mRNA destabilization effects were minimal by performing differential gene expression analysis (Figure S7A,B). Only a few differentially regulated genes were found, compared with thousands observed after long-term METTL3 knockout/knockdown (Batista et al., 2014; Geula et al., 2015; Ke et al., 2017; Yue et al., 2015). We then employed LeafCutter to perform intron-centric annotation-free differential splicing analysis (Li et al., 2018). The splicing events were sorted into 4 groups by graded significance, with Set1 as the most splicing changed group. The higher the significance cutoff, the higher was the proportion of differential splicing events that include or neighbor m6A peaks, with 72.5% in Set1 with the most significant threshold, and only 19% in Set4 with the lowest threshold (Figure 5B). Consistent with this, the m6A peak intensities and peak numbers overlapping Set1 splicing clusters were significantly higher than the remaining groups (Figure 5C). These analyses of the early response in the nascent transcriptome implies that METTL3 complex affects the inclusion of specific splicing elements by depositing m6A modifications at nearby splice sites.

**Figure 5.**
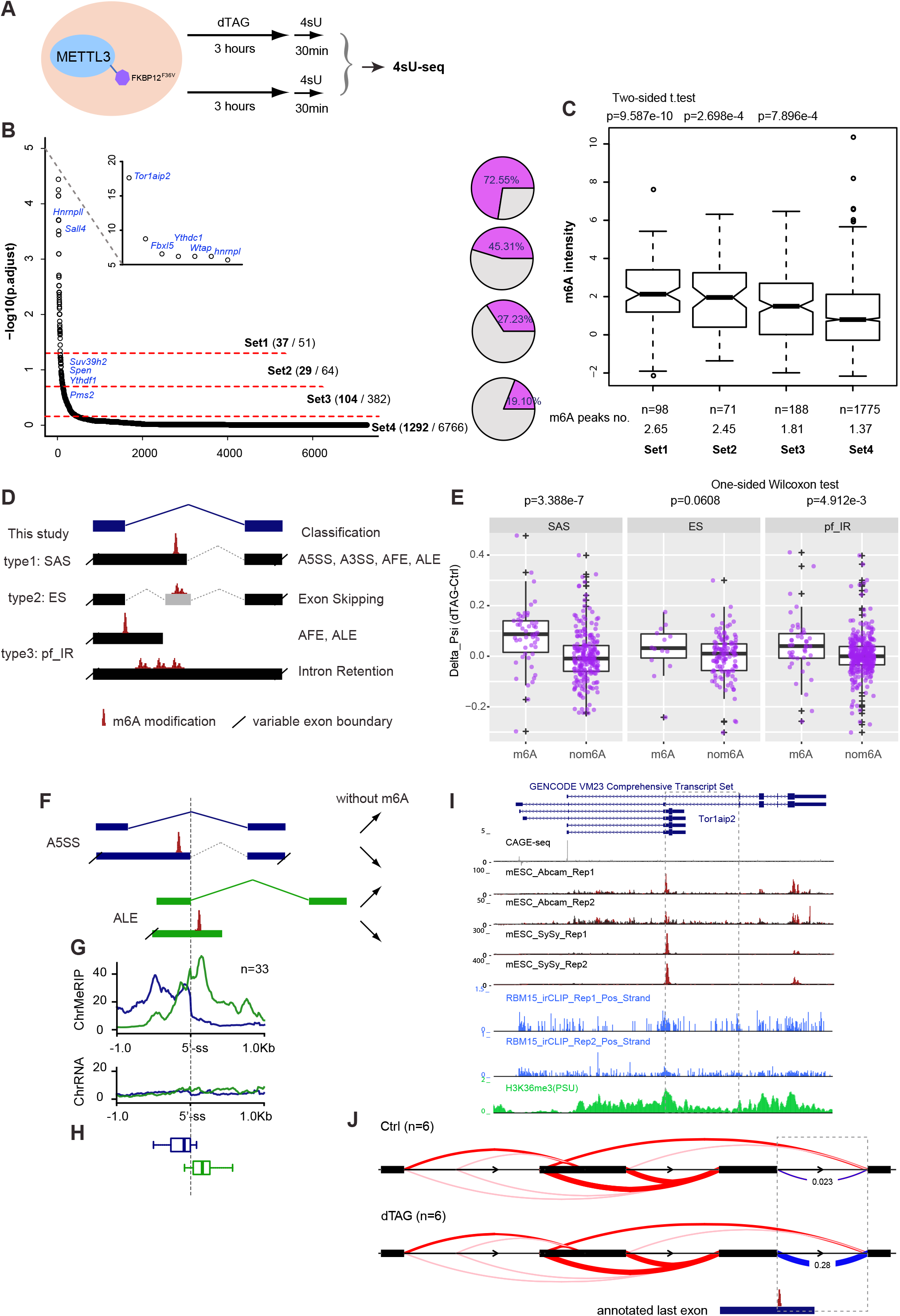
Rapid depletion of METTL3 causes m6A-targeted alternative splicing. (A) Schematic of 4sU-seq experimental design. (B) The output from intron-centric software LeafCutter ranks splicing clusters that change upon dTAG-13 treatment. Splicing clusters are ordered on the X axis according to their significance, which is plotted on the Y-axis (-log10(p.adjust)). Three different cut-offs were set to produce four groups graded by splicing significance. Selected genes are labelled. Pie charts (right) show the fraction of splicing cluster having or neighboring m6A modification. (C) Boxplots showing the m6A intensity distribution calculated from ChrMeRIP-seq. m6A peaks from high to low significance groups are ordered from left to right. P-values above boxes were calculated by a two-sided t-test for each group with respect to the non-splicing-change group (far right). The total peak number and average peak intensity for each splicing cluster are shown below. (D) Schematic showing types of splicing classified by type, with nomenclature used in this study (left) and canonical splicing classification (right). (E) Boxplots comparing the deltaPSI for each group (SAS, ES, and pf_IR). For each, splicing clusters with m6A modification located at the alternative intron/exon (left) are compared to splicing clusters from same group without m6A modification (right) as a batch-matched control. P-values shown above were calculated by one-sided Wilcoxon test. Positive deltaPSI indicates increased inclusion upon depletion of METTL3. (F) Splicing type A5SS and ALE. Right panel shows the direction of splicing changes upon functional m6A loss. (G-H) Aggregate m6A signals over the 5’-splice sites for A5SS and ALE (n=33 for both). Centre indicates the 5’-splice site, upstream and downstream 1kb are also included. Blue and green curves represent sites of A5SS and ALE, respectively. Bottom boxplots (H) denote the distance distribution towards the closest m6A peak summit. (I) Genome browser tracks for Tor1aip2 gene. From top to bottom, they denote CAGE-seq, MeRIP-seq (Abcam 2 replicates, SySy 2 replicates), Rbm15 irCLIP-seq (2 replicates), and H3K36me3 ChIP-seq. The dashed box depicts the most significantly changed splicing cluster in this study. (J) Sashimi plot of the Tor1aip2 splicing cluster. The dashed box is the same as (F), with deltaPSI calculated from LeafCutter. The annotated last exon containing m6A modification at the splicing site is shown below.

We next focused on characterizing alternative splicing defects caused by Mettl3 depletion using the aforementioned Set1-3 clusters. We grouped the intron-centric alternative splicing clusters from the LeafCutter output into three types: (I) Splice Alternative Site (SAS), (II) Exon Skipping (ES), (III) partial or full Intron Retention (pf_IR) (Figure 5D and Methods). To avoid batch effects, significant splicing changes occurring at sites without m6A modification from the same sample were chosen for comparison. To evaluate these different types, we always considered the splicing event that skips the alternative splicing element (Figure 5D and Methods) and measured splicing changes upon dTAG-13 treatment (deltaPSI) by changes to this longer splicing event. The deltaPSI of the longer splicing form is always reciprocal to changes of inclusion of the alternative splicing form that bears m6A modification (Figure 5D,E). This analysis shows that for all the described splicing types, the longer splicing forms are increased compared to controls following dTAG-13 treatment, most significantly for SAS and pf_IR types (Figure 5E).

When splicing direction was considered in the m6A-linked splicing events, we found that two thirds (33 out of 50) of SAS events have alternative 5’-splice site and three quarters (33 out of 43) of pf_IR events overlap with 5’-splice site, which agrees with the nascent m6A pattern that intronic m6A modifications are generally in proximity to 5’-splice sites. Intriguingly, although both SAS and pf_IR events are associated with intron/exon inclusion, we found that for SAS events, m6A near to an alternative 5’-splice site generally promotes splicing, whereas in pf_IR events, m6A located in the alternative last exon suppresses splicing (Figure 5F). With these observations in mind we sought to determine if m6A positional differences could account for the distinct splicing outputs. The distance between the 5’-splice site and the corresponding closest m6A peak summit was calculated accordingly. We found that generally m6A methylations promoting splicing reside in upstream of the 5’-splice site (median: 107nt), and m6A methylations suppressing splicing locate at downstream of the 5’-splice site (median: 140nt) (Figure 5G,H). A similar pattern was also seen using the RBM15 irCLIP data (Figure S7C). Together these findings illustrate how positioning of m6A in introns relative to the 5’-splice sites provides a basis for specifying distinct alternative splicing outcomes.

We went on to compare our findings with those from stable Mettl3 knockout mESCs (datasets from Ke et al. 2017 and Geula et al. 2015). Applying a similar pipeline, we found that only 25-34% of the splicing clusters include or neighbor our annotated m6A peaks, and this percentage decreases alongside the m6A peak number per splicing cluster from Set1 to Set4 (Figure S8A-F). Furthermore, the nascent splicing changes upon acute depletion of METTL3 barely overlapped with the splicing changes observed in the mature transcriptome from constitutive knockouts of the gene encoding Mettl3 (Geula et al., 2015; Ke et al., 2017) (Figure S8G), indicating that secondary effects on splicing are dominant in stable Mettl3 knockout mESCs. We did however find evidence that Mettl3 knockout has a small effect on m6A-containing cassette exon skipping, as reported by Xiao et al. (2016) (Figure S8E,F). This predominance of secondary effects is not surprising given that Mettl3 knockout affects a variety of RNA processing steps such as nuclear export (Roundtree et al., 2017b) and RNA decay (Ke et al., 2017; Liu et al., 2020; Wang et al., 2014).

Intron 3 of the Tor1aip2 gene was the most strongly affected differentially spliced gene in our acute METTL3 depletion experiment (Figure 5B), and interestingly also ranks among the top four in two previous studies (Geula et al., 2015; Ke et al., 2017) (Figure S8A,C). m6A modification renders the short isoform dominant under normal conditions, whereas the longer isoform (alternative last exon) becomes extensively induced, with loss of m6A significantly increasing the splicing at this junction (Figure 5I,J and S9A). Similar results reported to occur with conditional knockout of the nuclear m6A reader Ythdc1 in mESC (Liu et al., 2020) further demonstrate the role of m6A as opposed to other functions of the METTL3/14 complex in regulating splicing (Figure S9B). In summary, these results indicate that m6A can function to mediate inclusion of m6A-containing alternative introns/exons in the context of the nascent transcriptome.

### Auto-regulation of the m6A machinery by alternative splicing

Among thousands of m6A targets in mESCs, we intriguingly found that all of the m6A cytosolic reader genes Ythdf1/2/3 are site-specifically modified by m6A in their internal long CDS-coding exons (Figure S10C-E). The nuclear m6A reader YTH-containing protein Ythdc1 is also heavily methylated by m6A across exon11 and intron11 regions (Figure 1G). Accessory proteins of the core m6A heterodimer writer complex including Wtap, Cbll1, and Rbm15/15B are also extensively methylated, as well as two m6A erasers (Fto and Alkbh5) (Figure 6A and S10A,B). Additionally, Spen, an RRM and SPOC-domain-containing protein in the same family as RBM15 that has been implicated in m6A regulation (Dossin et al., 2020), is also heavily m6A methylated across exonic and intronic regions (Figure 1G). Together these results suggest the existence of feedback loops for regulating cellular mRNA m6A metabolism. Of the aforementioned examples, remarkably Wtap, Ythdc1, Ythdf1, and Spen all show m6A-dependent regulation of alternative splicing as an early response to m6A loss. The splicing changes occurring at these gene loci are of different types, Ythdc1 intron11 and Spen intron2 are SAS (alternative 5’-splice site), Wtap intron6 is pf_IR (alternative last exon) (Figure 6B,C), while Ythdf1 cassette exon is ES (exon skipping) (Figure 6D). Notably, splicing of Ythdc1 intron11 and Wtap intron6 are also significantly changed and rank as top hits in both stable knockout datasets (Geula et al., 2015; Ke et al., 2017) (Figure S8A,C).

**Figure 6.**
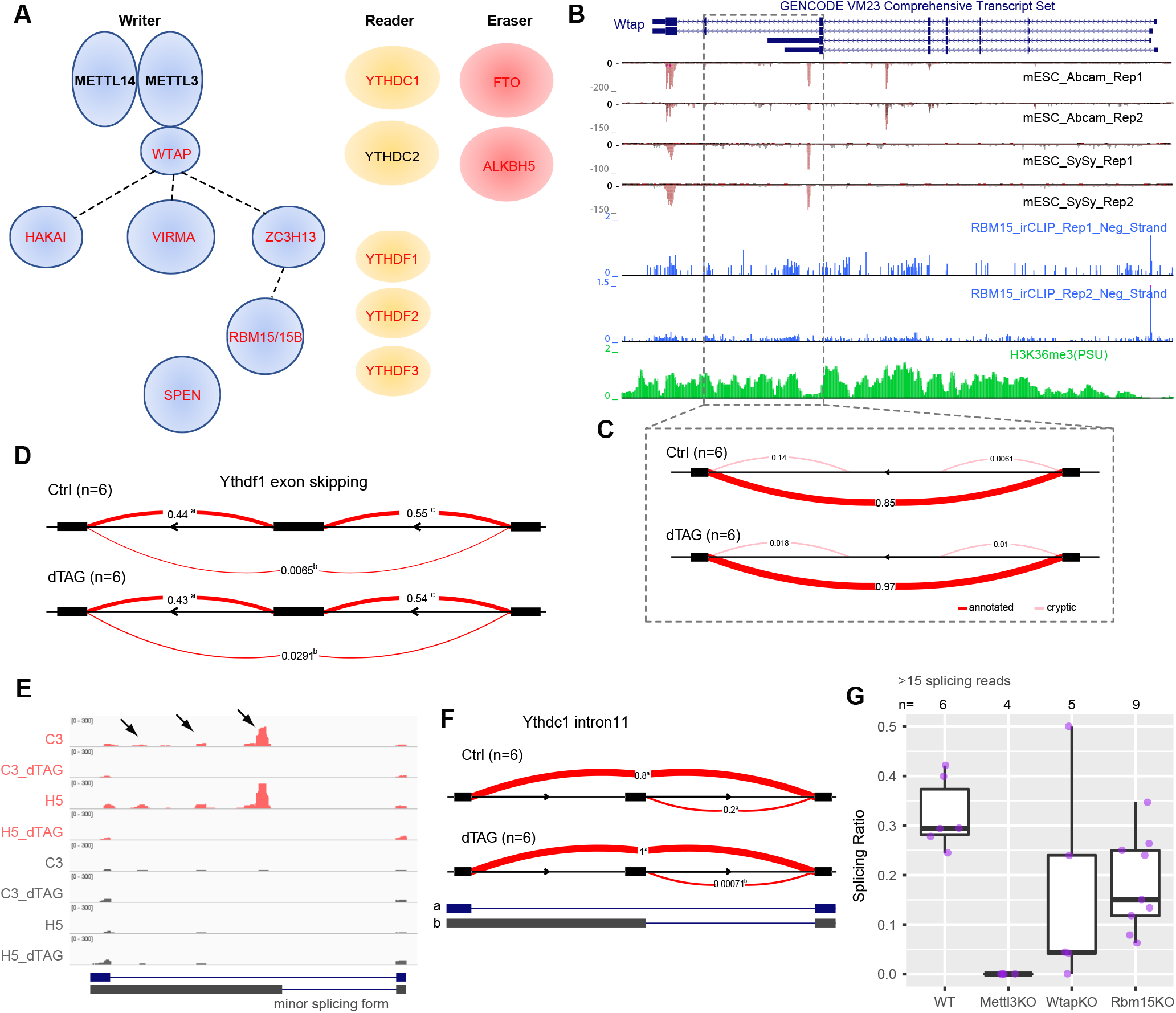
Splicing changes contribute to m6A self-regulation as an early consequence of acute m6A loss. (A) Schematic of m6A-related genes in writer, reader, and eraser complexes. Genes containing m6A modification are indicated by red text. Dashed lines in the writer complex indicate biochemically uncharacterized interactions. (B-C) Genome browser tracks (B) and Sashimi plot showing the detalPSI (C) for Wtap. Like the Tor1aip2 gene shown in Figure 5F-G, the splicing is of type ‘pf_IR’. (D) Sashimi plot showing the deltaPSI calculated from LeafCutter for Ythdf1 gene, ES type. (E) Genome Browser (IGV) tracks showing MeRIP-seq signal (top 4 tracks) and input signal (bottom 4 tracks) for Ythdc1 intron11. Arrowheads indicate m6A peaks located in the alternative intron part. Annotated splicing forms are shown below. (F) Sashimi plot showing changes for Ythdc1 intron11, which is of type ‘SAS’. (G) Boxplots showing the splicing choice score for the minor splicing form of Ythdc1 intron 11 from ChrRNA-seq datasets in which components of m6A writer complex were perturbed (Nesterova et al 2019). Samples were included only if more than 15 reads span the junction at Ythdc1 intron 11.

For these examples, a splicing choice score was calculated for the minor splicing forms (alternative 5’-splice site) (Methods). The splicing choice score for the Ythdc1 intron11 minor splicing junction is approximately 20-30% in wild-type cells, whereas it drops to nearly 0 in both acute and stable Mettl3 knockout cells (Figure 6E-G). This suggests that m6A modifications determine the inclusion of this minor splicing form. This is unlikely to be the consequence of RNA destabilization by m6A modification, which would result in the minor splicing form being overrepresented. When we explored ChrRNA-seq datasets generated from knockout of the accessory Mettl3/14 complex proteins Wtap and Rbm15 mESCs (Nesterova et al., 2019), we also observed significantly reduced representation of the minor splicing forms, although these effects were less drastic than in the Mettl3 knockout, probably due to the residual m6A levels in the accessory protein knockouts. Intriguingly, as it is the major splicing form of Ythdc1 that produces the main Ythdc1 protein-coding isoforms, loss of m6A modification at intron 11 (and thus the alternatively spliced short isoform) contributed to more efficient Ythdc1 transcript production. Accordingly, the Ythdc1 transcript (Figure S7A) and protein levels (Figure 4C) are higher in the acute METTL3 knockout mESCs. Taken together, splicing changes occurring as the immediate consequence of loss of m6A contribute to m6A self-regulation.

## Discussion

Our analysis reveals that in mESCs around 10% of m6A regions are located in introns, in broad agreement with a prior analysis of nascent RNA from HeLa cells (Ke et al., 2017). We observe preferential location of intronic m6A close to 5’-splice sites. Both intronic and exonic m6A regions show dependence on the METTL3/14 writer complex, as determined by acute depletion of METTL3 using the dTAG system. Our data further indicate that intronic m6A modifications are deposited through the same mechanisms as those reported to function in exons, RBM15 binding and H3K36me3-modified chromatin (Huang et al., 2019; Patil et al., 2016). Of note we observed preferential Rbm15 binding both around the stop codon/3’UTR, and at the start of transcripts, with only the latter correlating with the distribution of m6A. Similar binding profiles were observed in a prior study analyzing human RBM15/15B protein (Patil et al., 2016). The basis for preferred binding near the start of transcripts is currently unknown but may be linked to interaction of the RBM15 SPOC domain with the H3K4me3 methyltransferase SET1B that localizes to gene promoters (Coker et al., 2020; Lee and Skalnik, 2012). Accordingly, RBM15 binding close to the start of transcripts, which does not correlate with m6A levels, may have a distinct function.

A role for intronic m6A in regulation of splicing has been proposed previously, notably in relation to female specific *Sxl* splicing and sex determination in *Drosophila* (Haussmann et al., 2016; Lence et al., 2016). Consistent with this idea we observe preferential association of intronic m6A with alternatively spliced regions and moreover, following acute depletion of METTL3, widespread perturbation of splicing events in nascent RNA. Thus, we find that RNA m6A modifications are located in alternative introns/exons, including alternative 5’-splicing, alternative 3’-splicing, alternative last exons, and exon skipping isoforms, and that they can mediate inclusion of this alternative part in the nascent transcriptome. We further found that the location of m6A peaks relative to the 5’-splice site correlates with the splicing output. Our analysis thus extends previous models that either only covered exon skipping (Xiao et al., 2016) or focused on intron retention (Fish et al., 2019). Whilst we observed some overlap with splicing changes seen in prior studies, for example in the Tor1aip2 gene (Figure 5), our analysis detected many more instances of m6A-mediated differential splicing. The fact that our observations followed acute depletion of METTL3 suggests the changes are directly linked to METTL3 function rather than secondary long-term effects from perturbing the m6A system. A likely explanation for the relatively high number of aberrant splicing events that we detected is that we analysed splicing patterns in nascent RNA rather than processed mRNA. It follows that the affected introns/exons are underrepresented in processed mRNA samples, either because they are highly unstable or because they escape export and are retained in the nuclear chromatin fraction. Consistent with the former possibility, recent work has shown that m6A marks RNA transcribed from specific repeat sequence elements for degradation by the nuclear exosome (Liu et al., 2020).

We find evidence that genes encoding several subunits of m6A writer and reader complexes have m6A dependent splicing. Moreover, we observed that protein levels of some of these factors, for example YTHDC1, increase following acute depletion of METTL3, indicating that feedback mechanisms have evolved to regulate m6A dependent functions. In this specific case we infer that the m6A dependent splice form suppresses levels of the major protein coding splice variants, either as a result of altering ratios of translationally productive and non-productive mRNA or by a function for the non-productive transcript in transcription/translation in cis or in trans. It will be interesting in the future to further investigate this idea and to determine if other m6A dependent splicing events play a role regulating and/or fine tuning different biological pathways. The use of acute METTL3 depletion, in allowing discrimination of direct and indirect deficits, will be an important tool for any such studies. Of note, our finding that acute depletion of METTL3 leads to degradation of METTL14 to which it is stably bound, but not of other accessory proteins such as WTAP and RBM15 provides support to the suggestion that METTL3/14 forms a stable sub-complex distinct from the accessory proteins (Bokar et al., 1997; Knuckles et al., 2018).

In summary, we have defined the pattern of intronic m6A modification in the mESC nascent transcriptome and shown that intronic m6A mediates inclusion of alternative intron/exons, highlighting a potentially important level of gene regulation for the evolution and fine tuning of biological pathways.

## Acknowledgement

We are grateful to Nathanael Gray and James Bradner for sharing the dTAG plasmid and dTAG-13 reagents. We thank Julian Zagalak from Ule lab for help with RBM15 irCLIP, Tatyana Nesterova for generating the Xist inducible XX mESC cell line and general experimental assistance, Charlotte Capitanchik and Brockdorff lab members for suggestions and discussions, Huijuan Feng from Columbia University for critical reading of the manuscript, Amanda Williams from Oxford Zoology for NextSeq sequencing, Oxford Biochemistry IT support for computing server maintenance. Work in the Brockdorff lab is supported by the Wellcome Trust (grant no. 215513). The Francis Crick Institute receives its core funding from Cancer Research UK (FC001110), the UK Medical Research Council (FC001110), and the Wellcome Trust (FC001110).

## Author Contribution

GW and NB conceived and designed the project. GW optimized and performed the ChrMeRIP-seq. GP and JB initiated and optimized the RBM15 irCLIP protocol with the advice and reagents from JU. GP and GW performed the RBM15 irCLIP-seq in duplicates. GW and MA created the METTL3_FKBP12^F36V^ cell line. GW performed calibrated MeRIP-seq, 4sU-seq, and western blot with MA. HC adopted MeRIP-seq protocol. GW performed all the bioinformatics analysis, analyzed data, and generated figures. GW and NB wrote the manuscript with input and suggestions from co-authors. NB supervised the project and secured the funding.

## Materials and Methods

### Cell culture

All mouse embryonic stem cells (mESCs), except E14 mESCs, were grown in feeder-dependent conditions on gelatinized plates at 37°C in a 5% CO_2_ incubator. Mitomycin C-inactivated mouse fibroblasts were used as feeders. mESCs medium consists of Dulbecco’s Modified Eagle Medium (DMEM, ThermoFisher) supplemented with 10% foetal calf serum (Seralab), 2 mM L-glutamine (ThermoFisher), 1X non-essential amino acids (ThermoFisher), 50 μM *β*-mercaptoethanol (ThermoFisher), 50 g/mL penicillin/streptomycin (ThermoFisher), and 1mL of Leukemia Inhibitory Factor (LIF)-conditioned medium made in-house. E14 cells were grown feeder-free in the same conditions and medium as above. Xist expression was induced by the addition of 1–1.5 µg/mL doxycycline (Dox) (Sigma, D9891) for 24 hrs.

### Cell lines

E14 mouse embryonic stem cells (mESCs) were used for ChrMeRIP-seq. Hybrid (Cast/129S) XX mESCs having inducible Xist from Cast allele (Nesterova et al., 2019) were used to knockin FKBP12^F36V^ into METTL3 locus. The emGFP-PreScission-RBM15 cell line was derived from mouse XY 3E ES cells, containing rtTA integrated into the Rosa26 locus and random integration of Dox-inducible Xist transgene into chr17 (Coker et al., 2020; Tang et al., 2010). In these cells, the puromycin resistance cassette at Rosa26 locus was replaced with hygromycin resistance (Moindrot et al., 2015). Then, cells were transfected and screened for stable integration of the pTRE-emGFP-PreScission-RBM15 plasmid. Cells treated with 1μg/mL Dox for 24hr simultaneously induce Xist RNA and emGFP-PreScission-RBM15 protein expression.

### Molecular cloning and CRISPR-Cas9 mediated knockin

sgRNA targeting near Mettl3 stop-codon was designed by online tool CRISPOR (Haeussler et al., 2016) (http://crispor.tefor.net/) and oligos were synthesised from Invitrogen, then cloned into pSpCas9(BB)-2A-Puro (PX459) V2.0 (Addgene #62988) backbone following instructions. A donor vector was built by Gibson assembly (NEB) of homology arms (∼400bp) PCR amplified from genomic DNA and the FKBP12^F36V^ sequence amplified from plasmid pLEX_305-N-dTAG (Addgene # 91797) (Nabet et al., 2018). The Cas9-sgRNA-containing plasmid and donor vector were co-transfected at a molar ratio of 1:6 into XX mESCs cells on a 6-well plate using Lipofectamine 2000 according to the manufacturer’s protocol (ThermoFisher). Transfected cells were passaged at different densities into three Petri dishes with feeders, 24h after transfection. The next day, cells were subjected to puromycin selection (3.5 μg/mL) for 48h and then grown in regular mESC medium until mESC colonies were ready to be picked and expanded. Western blot analysis was used to screen colonies for FKBP12^F36V^ knock-in as this results in slower mobility of the METTL3-FKBP12^F36V^ protein compared to the wild-type protein in a SDS-PAGE gel and disappearance of the wild-type METTL3 band on western blot. Selected clones were further characterized by PCR of genomic DNA and Sanger sequencing to confirm correct knock-in and homozygosity. The sensitivity of selected METTL3-FKBP12^F36V^ clones to dTAG13 treatment was validated by western blot.

### Western blot

Total cell lysates were resolved on a polyacrylamide gel and transferred onto PVDF or nitrocellulose membrane by semi-dry transfer (15V for ∼50 min) or quick transfer. Membranes were blocked by incubating them for 1 hr at room temperature in 10 mL PBS, 0.1% Tween (TBST) with 5% w/v Marvell milk powder. Blots were incubated overnight at 4°C with the primary antibody, washed 4 times for 10 min with PBST and incubated for 1 hour with secondary antibody conjugated to horseradish peroxidase. After washing 5 times for 5 min with PBST, bands were visualised using ECL (GE Healthcare). TBP serves as loading control.

### Cell fractionation and Chromatin-associated RNA (ChrRNA) isolation

ChrRNAs were extracted from a confluent 15 cm dish of mESCs. Briefly, cells were trypsinised and washed twice in cold PBS, and then lysed on ice in RLB buffer [10 mM Tris pH 7.5, 10 mM KCl, 1.5 mM MgCl_2_, and 0.1% NP40] for 5min, and nuclei were purified by centrifugation through a sucrose cushion (24% sucrose in RLB). The supernatant was kept as cytosolic fraction. The nuclei pellet was resuspended in NUN1 [20 mM Tris pH 7.5, 75 mM NaCl, 0.5 mM EDTA, 50% glycerol], then lysed with NUN2 [20 mM Hepes pH 7.9, 300 mM, 7.5 mM MgCl_2_, 0.2 mM EDTA, 1 M Urea]. Samples were incubated for 15 minutes on ice with occasional vortexing, then centrifuged for 10min at 2800g to isolate the insoluble chromatin fraction. The upper phase was kept as nucleoplasm fraction. The insoluble chromatin pellet was resuspended in Trizol by passing multiple times through a 23-gauge needle. The disrupted chromatin pellet was either frozen in -80° C or freshly purified through standard Trizol/chloroform extraction followed by isopropanol precipitation. Samples were then treated with two rounds of Turbo DNaseI (AM1907, Life Technology) in order to eliminate any potential DNA contamination. Concentration and quality of RNA isolated from the chromatin pellet were determined by Nanodrop and Agilent RNA Bioanalyzer.

### ChrMeRIP-seq and Calibrated MeRIP-seq

MeRIP-seq was based on the method by (Dominissini et al., 2013) with minor modifications. Briefly, total RNAs or ChrRNAs were isolated from pre-plated ES cells according to the procedure above. RNA was fragmented by incubation for 6 min at 94°C in thin-walled PCR tubes with fragmentation buffer [100 mM Tris-HCl, 100 mM ZnCl_2_]. Fragmentation was quenched using stop buffer [200 mM EDTA, pH 8.0] and incubation on ice, before ensuring the correct size (∼100 bp) using RNA Bioanalyzer. Total RNAs isolated from Drosophila SG4 cells were also fragmented in parallel Eppendorf tubes. For ChrMeRIP (conventional MeRIP-seq on chromatin-associated RNA), approximately 50 ug ChrRNA were used. For calibrated MeRIP-seq using total RNAs isolated from dTAG13-treated and untreated control cells (C3 and H5, depicted in Figure 4), 300 ug fragmented (∼100nt) RNAs, supplemented with 30ug fragmented Drosophila total RNAs were mixed in m6AIP buffer. RNAs were incubated with 10 ug anti-m6A antibody (Synaptic Systems, 202 003; or Abcam #ab151230), RNasin (Promega), 2 mM VRC, 50 mM Tris, 750 mM NaCl and 5% Igepal CA-630 in DNA / RNA low-bind tubes for 2 hrs before m6A-containing RNA was isolated using 200 ul Protein A magnetic beads per IP (pre-blocked with BSA). After 2 hour incubation, extensive washing (1x IP buffer [10mM Tris-pH7.4, 150mM NaCl, 0.1%NP-40], 2x LowSalt buffer [50mM Tris-pH7.4, 50mM NaCl, 1mM EDTA, 1%NP-40, 0.1%SDS], 2x HighSalt buffer [50mM Tris-pH7.4, 1M NaCl, 1mM EDTA, 1% NP-40, 0.1%SDS], 1xIP buffer) were carried out to remove the unspecific binding. 6.7 mM m6A (Sigma) was used to elute RNA from the beads. Input and eluate samples were EtOH co-precipitated with Glycoblue, quantified and pooled as libraries generated using TruSeq Stranded total RNA LT sample prep (Abcam ChrMeRIP-seq experiment) or NEBNext® Ultra™ II Directional RNA Library Prep (SySy ChrMeRIP-seq) according to manufacturer’s instructions, but skipping the fragmentation step. 75bp single reads were obtained using Illumina NextSeq500.

### 4sU-RNA immunoprecipitation

4sU-RNA was generated and isolated essentially as described in (Pintacuda et al., 2017; Rabani et al., 2011). In detail, 4-thiouridine (4sU, Sigma, T4509) was dissolved in sterile PBS and stored at -20°C. 4sU was thawed just before use and added to the cells in the growing media at a concentration of 500 μM. METTL3-FKBP12^F36V^ expressing cells (C3 and H5 clones) were treated with or without dTAG13 (100nM) for 3 hours, then exposed to 4sU-suplemented medium for 30 minutes. Cell culture medium was rapidly removed from cells and 5 mL of Trizol reagent (Life Technologies) was added. Total RNAs were isolated and any potential DNA contamination was removed using Ambion DNA-free DNase Treatment kit (Life Technologies) according to the manufacturer’s instructions. For each μg of total RNA, 2 μL of Biotin-HPDP (Pierce, 50mg EZ-Link Biotin-HPDP), previously dissolved in DMF at a concentration of 1 mg/mL, and 1 μL of 10x Biotinylation buffer [100 mM Tris HCl pH 7.4, 10 mM EDTA], was added. The reaction was incubated with rotation for 15 min at 25°C. RNA was transferred to Phase Lock Gel Heavy Tubes (Eppendorf), and an equal volume of chloroform was added. After vigorously mixing, tubes were left incubating for 3 min at 25°C and then centrifuged at 13,000rpm for 5 min at 4°C. The upper phase was transferred to new Phase Lock Gel Heavy Tubes, and chloroform added again. After further centrifugation, the upper phase was transferred to a tube containing an equal volume of isopropanol and 1/10 volume of 5 M NaCl. After inversion, the tubes were centrifuged at 13,000 rpm for 20 min at 4°C. Supernatant was washed in 75% ethanol and resuspended in water. Biotinylated 4sU-RNA was recovered using the μMacs Streptavidin Kit (Miltenyi), with a modified protocol. Per μg of recovered biotinylated 4sU-RNA, 0.5 μL of streptavidin beads were added, in a total volume of 200 μL. Samples were incubated with rotation for 15 min at 25°C. μMacs columns supplied with the μMacs Streptavidin Kit were equilibrated in 1 mL of washing buffer [100 mM Tris HCl pH 7.5, 10 mM EDTA, 1 M NaCl, 0.1% Tween 20] at 65°C. Samples were added to the columns that were then washed 6 time with washing buffer, 3 times at 65°C and three times at 25°C. RNA was eluted in freshly prepared 100mM DTT. RNA was further purified using the RNeasy MinElute Cleanup kit (Qiagen) according to the manufacturer’s guidelines. 1 μL of 4sU-labelled RNA was quality-checked using the Agilent RNA 6000 Pico kit (Agilent technologies) according to the manufacturer’s instructions and run on a 2100 Bioanalyzer Instrument (Agilent).

### 4sU-seq

Approximately 750ng of 4su-labelled RNA were subjected to sequencing library preparation by TruSeq Stranded total RNA LT sample prep with ribo-zero gold (Cat. No. RS-122-2301) according to manufacturer’s instructions. Libraries were purified with AmPure X beads and quantified by Qubit. Equal moles of libraries with different barcodes were pooled together based on the KAPA quantification (Roche) and sequenced using Illumina NextSeq500 (FC-404-2002) in paired-end mode (2x 80).

### emGFP-RBM15 irCLIP

The emGFP-PreScission-RBM15 cell line was described above. irCLIP were performed as (Zarnegar et al., 2016) with few modifications. Briefly, cells were grown with feeders and harvested after 24h Dox treatment. 10 million mESCs were washed with ice-cold PBS irradiated once with 150 mJ/cm^2^ in a Stratalinked 2400 at UVC 254 nm on ice. Snap frozen pellets were stored at -80°C until use. Cell pellets were resuspended in 1 ml lysis buffer [50 mM Tris-HCl, pH 7.4, 100 mM NaCl, 1% Igepal CA-630, 0.1% SDS, 0.5% sodium deoxycholate] supplemented with protease inhibitors, and transferred to a fresh 1.5 mL tube. Sonication was performed on a Bioruptor (Diagencode) for 10 cycles with alternating 30 secs on/ off at low intensity, and followed by RNase I (Thermo Scientific, EN0602) and Turbo DNase (Ambion, AM2238) treatment. Pre-washed antibody-conjugated beads were directly added to the RNase-treated lysate and rotated on a wheel overnight at 4°C. After discarding the supernatant, the beads were washed 2x with high-salt wash buffer [50 mM Tris-HCl, pH 7.4, 1 M NaCl, 1 mM EDTA, 1% Igepal CA-630, 01% SDS, 0.5% sodium deoxycholate]. 3’-end RNA dephosphorylation was performed on beads with PNK (NEB M0201L) for 20 min at 37°C. Beads were further washed with 1x PNK wash buffer [20 mM Tris-HCl, pH 7.4, 10 mM MgCl_2_, 0.2% Tween-20], 1x high-salt wash buffer, and 1x PNK wash buffer. Washed beads were subjected to RNA ligation with pre-adenylated infrared adaptor L3-IP-App (/5Phos/AG ATC GGA AGA GCG GTT CAG AAA AAA AAA AAA /iAzideN/AA AAA AAA AAA A/3Bio/, 1 µM) together with RNA ligase (M0204, NEB), and incubated overnight at 16°C in a thermomixer shaking at 1100 rpm. Beads were subsequently washed with1x PNK wash buffer, 1x high-salt wash buffer, 2x PNK wash buffer, and finally supplemented with 1x NuPAGE loading buffer. Supernatants were carefully collected and loaded on a 4-12% NuPAGE Bis-Tris gel (Invitrogen) according to the manufacturer’s instructions. The gel was run at 180V until dye front reached the bottom. The protein-RNA-L3IR complexes from the gel were transferred to a Protan BA85 Nitrocellulose Membrane (Whatman) using the Novex wet transfer apparatus according to the manufacturer’s instructions (Invitrogen; transfer for 2 hrs at 30 V). The entire lane above the expected molecular size of the protein of interest (emGFP-PreScission-RBM15) was cut from the membrane, and mixed with proteinase K (Roche, 03115828001). RNAs were isolated by Phenol:Chloroform:Isoamyl Alcohol (Sigma P3803) using Phase Lock Gel Heavy tube (713-2536, VWR). The isolated RNAs were reverse transcribed by SuperScript IV with barcode primer (/5Phos/ WWW GTGGA NNNN AGATC GGAAG AGCGT CGTGAT /iSp18/ GGATCC /iSp18/ TACTG AACCG C). cDNA products were biotin captured with MyOne C1 SA-dynabeads, eluted, and subsequently circularised with CircLigase II (Epicentre). PCR amplification was obtained after 18-21 cycles. The amplified irCLIP libraries were purified using AmPure XP beads (Beckman Coulter) and subjected to size selection from TBE gel by cutting a 150-500 nt smear band. Library concentrations were quantified by Qubit and KAPA quantification kit (Roche), quality-checked by DNA Bioanalyzer (Agilent), and sequenced on an Illumina NextSeq500 machine using 75bp single end sequencing.

### RNA m6A modification peak calling and confidence group classification

We performed peak calling on duplicate m6A IP and input alignment bam files with MACS2 (2.1.1) tool (Zhang et al., 2008). SySy and Abcam ChrMeRIP-seq data were analyzed separately. Nascent transcriptome size was calculated from UCSC genome browser and used as genome size. The key parameters are *(-q 0*.*05 --nomodel --extsize 100 --call-summits*) in addition to the genome size (*gsize*) 2.4e8 for ChrMeRIP and 1.05e8 for standard MeRIP-seq (Dominissini et al., 2013). The strand-specific bigwig files were generated from RNA-seq by BedTools (Quinlan and Hall, 2010) and peak strands were determined by calculating the strand-specific fold change log2(IP/Input) with UCSC utility *bigWigAverageOverBed*. Direction ambiguous peaks were removed from the analysis. The border of m6A peaks were further refined according to the summits output from MACS2. Given that RNAs were fragmented to a size slightly longer than 100nt for MeRIP-seq, we shrunk the peak size by only keeping the 125nt on both side of summits if the called m6A peak is larger than 250nt. For the subpeaks within broader peaks, we treated them as individual peak if they are 250nt further away. Otherwise, the border of these 2 subpeaks were determined as middle site between their summits. The custom scripts used for these refinements were also attached. This analysis resulted in 10,749 and 11,726 peaks for SySy and Abcam antibody respectively. We further defined the high confident m6A set (ConfGroup1, n=5277) if peak appears from 2 antibodies, medium confident m6A set (ConfGroup2, n=5472) that were detected by SySy antibody and also has signal from Abcam antibody but below the cutoff to be called peaks by MACS2 (Zhang et al., 2008), and low confidence m6A set (ConfGroup3, n=6319) that were only detected by Abcam antibody and almost no signal from SySy antibody. Motifs for each group were searched and analysed by HOMER program (Heinz et al., 2010).

### Intronic patterns of m6A methylation

Classification of peak summits (single nucleotides) were intersected with the selected MaxORF_LongestNcRNA GENCODE vM24 isoform (*intersectBed -a summits*.*bed -b \**.*GeneBed -s -wo*). Constitutive or alternative intron location was determined by whether the peak summit locates in intron for all the annotated comprehensive GENCODE vM24 isoforms or subsets of the isoforms respectively. PhastCons scores for multiple alignments of 59 vertebrate genomes to the mouse genome (phastCon60way) from UCSC were used (Pollard et al., 2010). For calculating conservation and GC contents, control region for this intronic m6A methylation was chosen by randomly selecting the size-matched region from the same intron. For the relative intron length and position, random intron from the same transcript was chosen. Plots were generated by R package ggplot2. Scripts used for this analysis were attached.

### Calibrated MeRIP-seq analysis

The raw reads from m6A immunoprecipitation were mapped to mouse (mm10) and Drosophila (dm6) concatenated genome by STAR (v2.5.2b) (Dobin et al., 2013). The resulted alignment files (bam) were then split into mm10 and dm6 alignment files according to the chromosome name. The read counts mapping to dm6 were counted (Li et al., 2009) and used for subsequent normalization (Table S1). The raw reads from input samples were mapped to mm10 genome by STAR (v2.5.2b) (Dobin et al., 2013). Alignment files for m6AIP and input samples were also normalized in a conventional manner by 10 million mapped reads. The strand-specific bigwig files were generated and visualized by IGV (Robinson et al., 2011).

### RNA m6A peak intensity analysis

MeRIP-seq reads were split into positive and negative strands (Li et al., 2009), and bigwig files were generated accordingly. Ten million mapped reads / library was used to perform normalization in ChrMeRIP and conventional MeRIP-seq data, except calibrated MeRIP-seq analysis that was normalized to the sequenced *Drosophila* RNA reads (Table S1). m6A Peak intensity was calculated as log2 RPKM_IP/RPKM_input. If this peak intensity is close to 0 or less than 0, it indicates that no m6A enrichment. m6A peaks were further classified into peaks close to transcript start site (TSS) as determined by deep -sequenced nuclear CAGE libraries from E14 mESCs (GSE148382) (Wei et al, 2020), peaks distal to TSS (genebody). Genebody m6A peaks were further grouped into intronic peaks, intergenic peaks. Peaks closer to TSS may be potential m6Am peaks because m6A antibody cannot distinguish m6A and m6Am.

### RNAmpp plot

RNA meta-profile plot (RNAmpp) scripts were written for this study. Step1, isoform selection (RNAmpp_prep.sh). Gene annotation were downloaded from GENCODE or UCSC genome browser as GTF format. The representative isoform can be chosen by 3 ways, (1) the random one (2) the one with maximum open reading frame (ORF) or the longest one if equal ORF exists; Longest lncRNA (3) the user-defined custom one. Step2, relative position calculation (RNAmpp_stat.sh). The single strand-specific nucleotide position (Bed-format file) called from m6A-seq or irCLIP-seq etc were used as input to find the intersected gene in a strand-specific manner with intersectBed (-s) from Bedtools (Quinlan and Hall, 2010), then iterated the introns or exons of overlapped genes to calculate the relative position to the feature start (the start position of the overlapped features i.e. 5’UTR, CDS, 3’UTR for protein-coding gene, transcript start sites for long ncRNA, and introns for all). Given that 5’UTR, CDS, and 3’UTR have varied lengths for genes, the averaged length for all the 5’UTR were calculated, so did CDS and 3’UTR. To account for these length differences, the common width (bin_size) represents similar length were searched based on the average length of 5’-UTR (221.23nt), CDS (1638.32nt), and 3’-UTR (1252.16nt), calculated from GENCODE vM24 annotation. This allows direct comparison among different regions from mRNA. For intron, each intron was split into 40 bins with equal size, and relative position for each query site was calculated as intron 5’-start. The pie chart and meta profile was generated by R packages. The dashed lines in meta profile denote the CDS start and CDS stop. The piecharts have two levels, level 1 shows the fraction of exon, intron, and intergenic, while level 2 further shows the protein coding fraction of exon and intron. RNAmpp, implemented in Python and R with dependency, is publicly available in github.

### irCLIP-seq analysis

The irCLIP adaptors (sequence) from single-end sequencing reads were first cut by cutadapt (v1.12) (Martin, 2011), and then collapsed by CTK toolkit to remove PCR duplication (Shah et al., 2017). The barcodes with UMI-containing 12 letters were moved to FASTQ header lines. The resulted single-end fastq reads, longer than 15nt, were mapped to mm10 genome by STAR (v2.5.2b) (Dobin et al., 2013) with key parameters (*--outSAMattributes All -- outFilterMultimapNmax 1 --outFilterMismatchNmax 2 --alignEndsType EndToEnd -- seedSearchStartLmax 15 --outWigType bedGraph read1_5p*). This setting is because RT stops were generated by reverse transcriptase around the UV crosslinked sites or crosslinking induced truncation sties (CITS), and RT-stops were main output when analysing the irCLIP-seq reads. The single nucleotide was considered as a CITS if there are no less than 3 stops (>=3) from independent strand-specific reads. In addition, crosslinking induced mutation sites (CIMS), particularly (deletion site) were also calculated. The single nucleotide was considered as a CIMS if this position has more than 2 deletions among total covered reads and the mutation rate is [0.01, 0.5], in order to avoid the potential contamination from the indels. Given that most of the RNA-binding proteins has 5-mer consensus motif, called CITSs or CIMSs from biological replicate 1 were kept if they were also called and within 5 nucleotides distance in biological replicate 2 (*closestBed -a CITS_Rep1*.*bed -b CITS_Rep2*.*bed -s -d* | *awk ‘$13<=5’*). The distribution of CITS and CIMS were analysed by the RNAmpp pipeline, described as above. Motifs around these CITSs or CIMSs were searched by Homer with key parameter (*findMotifsGenome*.*pl CITS*.*bed mm10 MotifOutput -rna*).

### RBM15 binding, m6A peak, and H3K36me3 modification

RBM15 CITS (>=3) called in 2 biological replicates were intersected with m6A peaks from different confident groups. Due to the strand-specificity, the closest distance between RBM15 binding and m6A peak will be minus if RBM15 CITS located upstream of m6A peak (*closestBed - a Rbm15_CITS*.*bed -b m6A_peaks*.*bed -s -D a*) (Quinlan and Hall, 2010). Strong Rbm15-binding group was defined by no more than 1000nt distance between Rbm15 binding and m6A peak. The density of Rbm15 binding sites (CTIS>=3) 1000bp flanking exonic m6A peaks, intronic m6A peaks, and intergenic m6A peaks were plotted. Bigwig files for H3K36me3 ChIP-seq data retrieved from ENCODE was generated in 10bp bin size. The metagene profile and heatmap for RBM15-binding group and other groups were generated by deeptools (v3.4.3) (Ramirez et al., 2016). The matrix generating from negative strand bigwig files are reversed and combined with the one from positive strand, and with a brief modification to average two replicates, then subjected to plotHeatmap tool (Ramirez et al., 2016).

### Differentially expressed genes analysis

4sU-seq were mapped to mm10 genome by STAR (v2.5.2b) (Dobin et al., 2013) allowing multiple mapping (*--outFilterMultimapNmax 100 --outFilterMismatchNmax 5 --alignEndsType EndToEnd --winAnchorMultimapNmax 100*). TEtranscripts (Jin et al., 2015) was used to detect the differentially expressed genes and repeat elements.

### Alternative splicing analysis

4sU-seq were mapped to mm10 genome by STAR (v2.5.2b) (Dobin et al., 2013) with following key parameters (*--twopassMode Basic --outSAMstrandField intronMotif --outSAMattributes All -- outFilterMultimapNmax 1 --outFilterMismatchNoverReadLmax 0*.*06 --alignEndsType EndToEnd*). For intron-centric differential splicing analysis, the leafcutter package (v0.2.7) (Li et al., 2018) was used to quantify intron usage and identify differentially spliced intron clusters between two conditions (dTAG13 treated and non-treated control, or wild-type versus Mettl3 knockout). Only splice junctions supported by uniquely mapped reads were used. The analysis followed the differential splicing documents from the package, except the split reads number for intron cluster. For dATG Mettl3 experiment, at least 30 split reads (6 replicates) are required but 25 for the published Mettl3 knockout datasets (GSE86336 from Ke et al 2017, GSE61997 from Geula et al 2015). The splicing clusters were ranked and 3 different thresholds were set to generate differential splicing level (For dTAG Mettl3 experiment, q<0.05, q<0.2, p<0.05 were chosen; For Mettl3 knockout datasets, q<0.01, q<0.05, p<0.05 were chosen. Here q is adjusted p-value), given that the RNA spliced reads coverages are different. If the splicing cluster has one (or multiple) m6A peak(s) from any confident group overlap or within 500bp distance, it was considered as *cis*-m6A-regulated cluster, otherwise other cluster. Sashimi plots with deltaPSI were generated from leafcutter package (Li et al., 2018). Peak intensity from each m6A peak was calculated as above and average m6A peak number was calculated from each cluster. The splicing choice score for alternative 5’-splice site was defined by splicing reads covering the minor splicing junction divided by the total splicing reads covering both minor and major splicing junction in this intron.

Three splicing types were described in this study based on the intron-centric analysis from leafcutter output, which is slightly different from the actual classification which has A5SS, A3SS, IR, ES, MXE, AFE, and ALE types. Here we described 3 splicing types: SAS, ES, and pf_IR (Figure 5). SAS (splicing alternative site) consists of two or more alternative splicing sites in the cluster and share either the start or the end coordinate. This type covers most of alternative 5’-splice site and alternative 3’-splice site (A5SS or A3SS), as well as alternative first exon (AFE) or alternative last exon (ALE) in some cases. ES are same as exon skipping model, where cassette exon either inclusive or exclusive in the spliced product. Partial or full intron retention (pf_IR) refers to the splicing junction partially overlaps with the first (AFE) or last exon (ALE) partially or fully inside of any exon, indicating that partial or full intron is included in the splicing product. This splicing type analysis were only focused on the clusters being identified as significant splicing change. To avoid batch effects, we did the comparisons for the same splicing types from same experiment with and without m6A modifications in the alternative exon/intron part (Figure 5). The distance between 5’-splice site and m6A peak summit was calculated using Bedtools (Quinlan and Hall, 2010) (*closestBed -a 5’-splice_site*.*bed -b m6A_peak_summit*.*bed -s -D a*).

## DATA AND CODE AVAILABILITY

High-throughput sequencing data (ChrMeRIP-seq, RBM15 irCLIP, Calibrated MeRIP-seq, and 4sU-seq) generated in this study have been deposited in Gene Expression Omnibus (GEO) under GSE154709. UCSC Genome Browser view of the ChrMeRIP-seq, RBM15 irCLIP-seq, and CAGE-seq (GSE148382) can also be accessed (http://genome-euro.ucsc.edu/s/Guifeng/ChrMeRIP). Scripts used in this study can be found (https://github.com/guifengwei/Nascent_m6A_Scripts), RNAmpp analysis (https://github.com/guifengwei/RNAmpp).

